# Nanopore adaptive sampling of a metagenomic sample derived from a human monkeypox case

**DOI:** 10.1101/2023.03.21.533647

**Authors:** C Hewel, H Schmidt, S Runkel, W Kohnen, S Schweiger-Seemann, A Michel, S-E Bikar, B Plachter, T Hankeln, M Linke, S Gerber

## Abstract

In 2022, a series of human monkeypox cases in multiple countries led to the largest and most widespread outbreak outside the known endemic areas. Genomic surveillance is of utmost importance to control such outbreaks. To this end, we performed Nanopore Whole Genome Sequencing of a local monkeypox sample on a PromethION 24. Adaptive sampling was applied for *in silico* depletion of the human host genome, allowing for the enrichment of low abundance viral DNA without *a priori* knowledge of sample composition. Nanopore sequencing allowed for high viral genome coverage, tracking of sample composition during sequencing, strain determination, and preliminary assessment of mutational pattern. Nanopore sequencing is a highly versatile method to characterize a virus in real-time without pre-sequencing target enrichment.

## Introduction

Monkeypox is a zoonotic disease caused by the enveloped monkeypox virus (MPXV) which contains double-stranded DNA with a genome size of 197 kb harboring around 200 genes. MPXV belongs to the *Orthopoxvirus* genus of the Poxviridae family. Three phylogenetically distinct clades of MPXV are known (I, IIa, IIb). MPXV of the 2022 outbreak belong to Clade IIb (Gigante et al. 2022), with a single origin potentially in the endemic region of Nigeria (Isidro et al. 2022). So far, more than 86,000 diagnosed cases and over 100 fatalities in more than 100 locations were counted in the 2022 MPXV outbreak (Mathieu et al. 2022). The natural reservoir of this most recent outbreak remains unknown, but MPXV is frequently found among rodents in Africa, even in 120-year-old historic specimen (Tiee et al. 2018). Human-to-human transmission of MPXV is most prevalent in Clade IIb and occurs primarily via direct contact with infected skin lesions and/or other body fluids and recently contaminated objects (Fleischauer et al. 2005, Hutson et al. 2013). The primary infection of the 2022 outbreak had a mean incubation time of roughly 9 days (UKHSA, 2022). Classical symptoms of the following prodromal phase (1-4 days) include fever, headache, fatigue, and, less often, lymphadenopathy. Lymphadenopathy is a distinctive clinical feature of monkeypox compared to other diseases that may initially appear similar (chickenpox, measles, smallpox). During the eruptive phase (14-28 days), skin lesions appear through several stages: macules, papules, vesicles, and finally pustules. The case fatality ratio of monkeypox has historically ranged from 0 to 11 % in the general population and has been higher among young children. In recent times, the case fatality ratio has been around 3–6% (WHO, 2022). Epidemiological and clinical features of the current outbreak indicate it is sexually transmitted, which has not been observed before (Bragazzi et al. 2022). The cessation of smallpox vaccinations, providing some cross-protection against monkeypox, has been discussed as one factor that has led to significantly increased case numbers over the last 13 years, cumulating in the outbreak of 2022. Over the same period, the median age at presentation has increased from 4 to 31 years (Petersen et al. 2019, Bunge et al. 2022). Taken together, monkeypox is gradually evolving to become of global public health relevance. Surveillance and detection programs are essential tools for understanding the continuously changing epidemiology of this resurging disease.

Integrating high-throughput sequencing metagenomic workflows into pathogen diagnostics has the power to disrupt traditional procedures for infectious diseases since turnaround time is dramatically reduced by waiving the cultivation of suspected pathogens. Moreover, it provides higher taxonomical resolution of microorganism kingdoms, even of those species that cannot be cultivated (Simner et al. 2018). PCR has become a gold standard for detection of viruses. Highly specific PCR assays for differentiating closely related viral species are increasingly difficult to maintain due to the high mutability of a viral genome and the entirety of human-infecting viruses are yet to be revealed. The applicability of clinical metagenomics has been evaluated in viral pathogen detection, demonstrating high potential benefits here as well (Kiselev et al. 2020). Despite the immense potential of high-throughput sequencing metagenomic workflows, there are also crucial limitations. Highly problematic is the high host-to-pathogen nucleic acid fraction in clinical samples with extreme values of more than 99% (Marquet et al. 2022, Martin et al. 2022). This unfavorable ratio can be overcome by complex and time-consuming laboratory methods, like differential lysis or enzymatic and immunomagnetic protocols. These solutions in turn can disturb the metagenomic composition of the sample (Horz et al. 2008, Liu et al. 2016, Ferretti et al. 2017, Oechslin et al. 2018). The limited read length of the current sequencing platforms also hampers the assignment of important genes (e.g., antimicrobial resistance) to the respective species and the alignment of reads to repetitive regions in order to cover the whole genome of pathogens (Yang et al. 2022). Nanopore sequencing bears the potential to overcome these limitations (Liu et al 2022). Due to its unrivaled read length, it can span complex genomic regions and resolve them unambiguously. Furthermore, it is technically possible to reverse the voltage across nanopores to enable selective sequencing of target fragments, based on real-time assessment of a small initial part of their sequence. This strategy, known as adaptive sampling (Loose et al. 2016), enables depletion of unwanted host-derived nucleic acid fragments (Cheng et al. 2022) and avoids rather complex wet lab sample purifications. The high efficiency of human host depletion in clinical samples via this approach has been shown by Marquet et al. (2022). Without changing the microbial composition, the authors showed a 1.7-fold increase in sequencing depth for samples with high human DNA contamination. While working with relatively short reads because of a PCR-based sequencing library preparation, the authors outlined that adaptive sampling’s decision-making would be more efficient with longer reads, further increasing overall sequencing depth. Using a PCR-free library preparation of a clinical MPXV sample, we aimed here to evaluate the potential of adaptive sampling towards a clinical routine in diagnosing culture-free metagenomic samples.

## Materials and Methods

### Sample acquisition, DNA isolation and qPCR testing

DNA extraction of a MPXV sample was performed on a single pustular lesion using the QIAamp DNA Mini kit, according to the manufacturer’s instructions. Extracted DNA was quantified using a Qubit dsDNA HS Assay (Invitrogen) following the manufacturer’s protocol and stored at −20°C until use. Initial testing for MPXV nucleic acid via qPCR analysis was positive (Ct value: 21) and performed on a NeuMoDx 288 Molecular System (Molecular Systems, Ann Arbor, USA).

### Metagenomic sequencing and bioinformatic processing

Ligation sequencing libraries were prepared from 1000 ng DNA using the Ligation Sequencing kit SQK-LSK110 (ONT, Oxford Nanopore Technologies Ltd., Oxford, UK) following the manufacturer’s protocol. The clean-up step after adapter ligation was intended to size-select fragments and was done with Short Fragment Buffer. The library was loaded on a single R9.4.1 (FLO-PRO002) flow cell and sequenced on a PromethION 24 device within 72 h. MinKNOW (v22.12.05) was used to supervise the initial sequencing run, including adaptive sampling on half a flow cell with depletion of human DNA by setting human genome build hg38 as input reference. NanoComp (v1.12.0; https://github.com/wdecoster/nanocomp) was used to determine effectiveness of the adaptive sequencing by comparing the respective subsets of the *sequencing_summary.txt file. After run completion, samples were re-basecalled and aligned to MPXV reference genome MT903344, while filtering out non-mapping reads using Guppy (v6.1.5) with the SUP model on a NVIDIA A100 DGX-station. All fastq reads were used as input for final species determination using Centrifuge (v 1.0.4-beta with the default h+p+v+c index) and visualised via Pavian (v1.0; https://github.com/fbreitwieser/pavian). The filtered reads were further used to obtain read quality metrics via SeqFailR (https://github.com/cdelahaye/SeqFaiLR) and NanoComp. Moreover, the filtered reads were re-formatted using bam2fq and used as input for the wf-mpx (v0.0.6; https://github.com/epi2me-labs/wf-mpx) pipeline for initial assessment and generation of a draft consensus with ON563414, an Oxford Nanopore-only assembly, as reference. This assembly overcomes limitations of short-read-only derived assemblies in terms of larger genomic rearrangements. Initial lineage assignment was performed via NextClade (v2.12.0; https://clades.nextstrain.org).

## Results

### Efficiency of adaptive sampling and lineage assignment of MPXV genome

In order to establish third generation metagenomic sequencing on a PromethION in our surveillance sequencing consortium and to characterise a monkeypox sample from the current outbreak, material from a skin lesion swap was prepared and loaded on a single R9 flow cell on a P24. During the run, adaptive sampling was activated for 1/2 half of the flow cell.

After 72 h of sequencing, 72.31 Gb of data were generated (see Fig. 1) consisting of 83.04 M reads. As evaluated by MinKNOW, N50 for a completed read was at 1.49 kb.

**Fig. 1:**
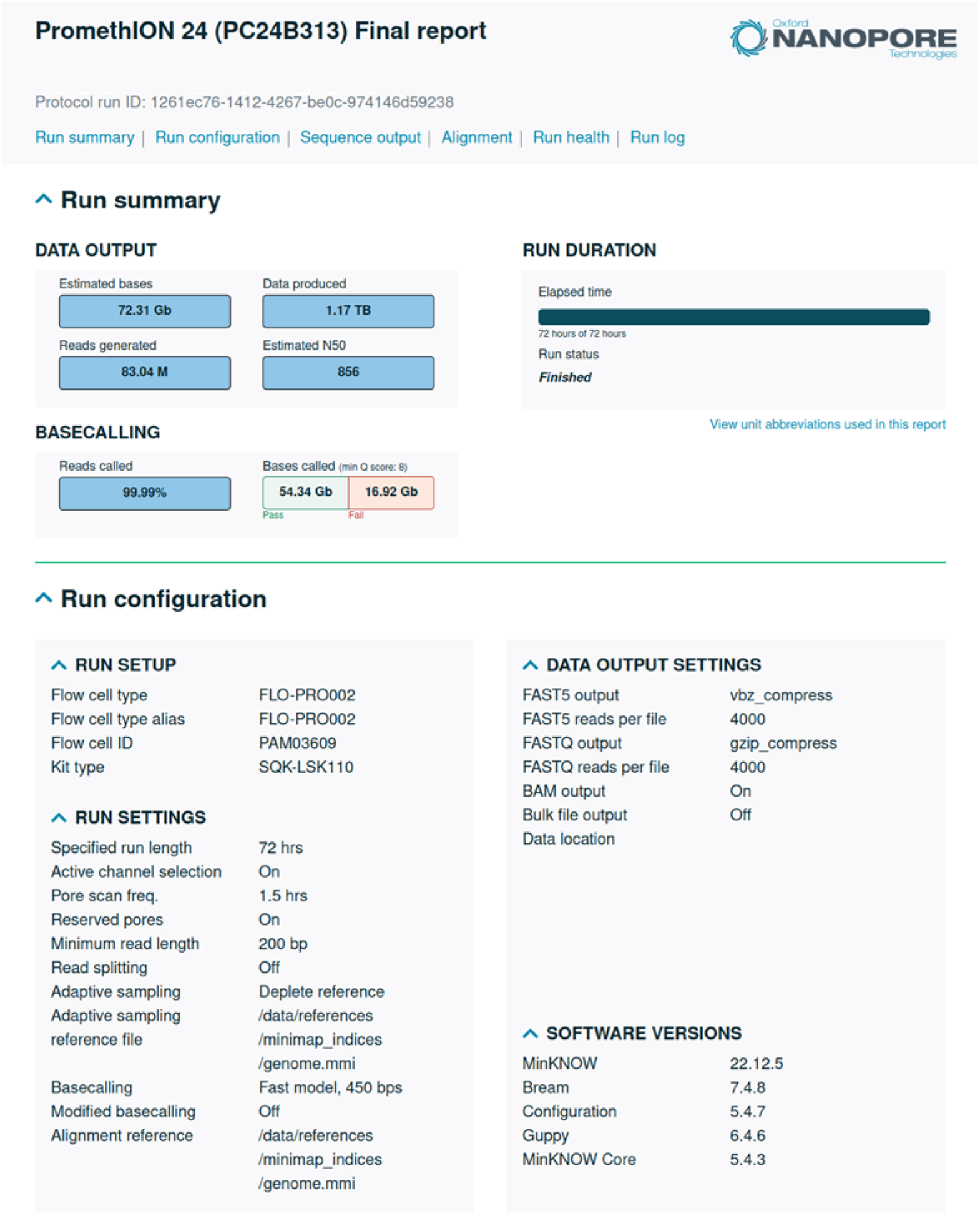
Excerpt from the run report by MinKNOW

The sequence quality and general error pattern was assessed with SeqFaiLR (see Fig. 2). Particularly in homopolymeric regions, errors become frequent and deletions start to become the predominant type of artifacts. Moreover, accuracy decreases with read length, dropping well below 50 % for longer homopolymer stretches.

**Fig. 2:**
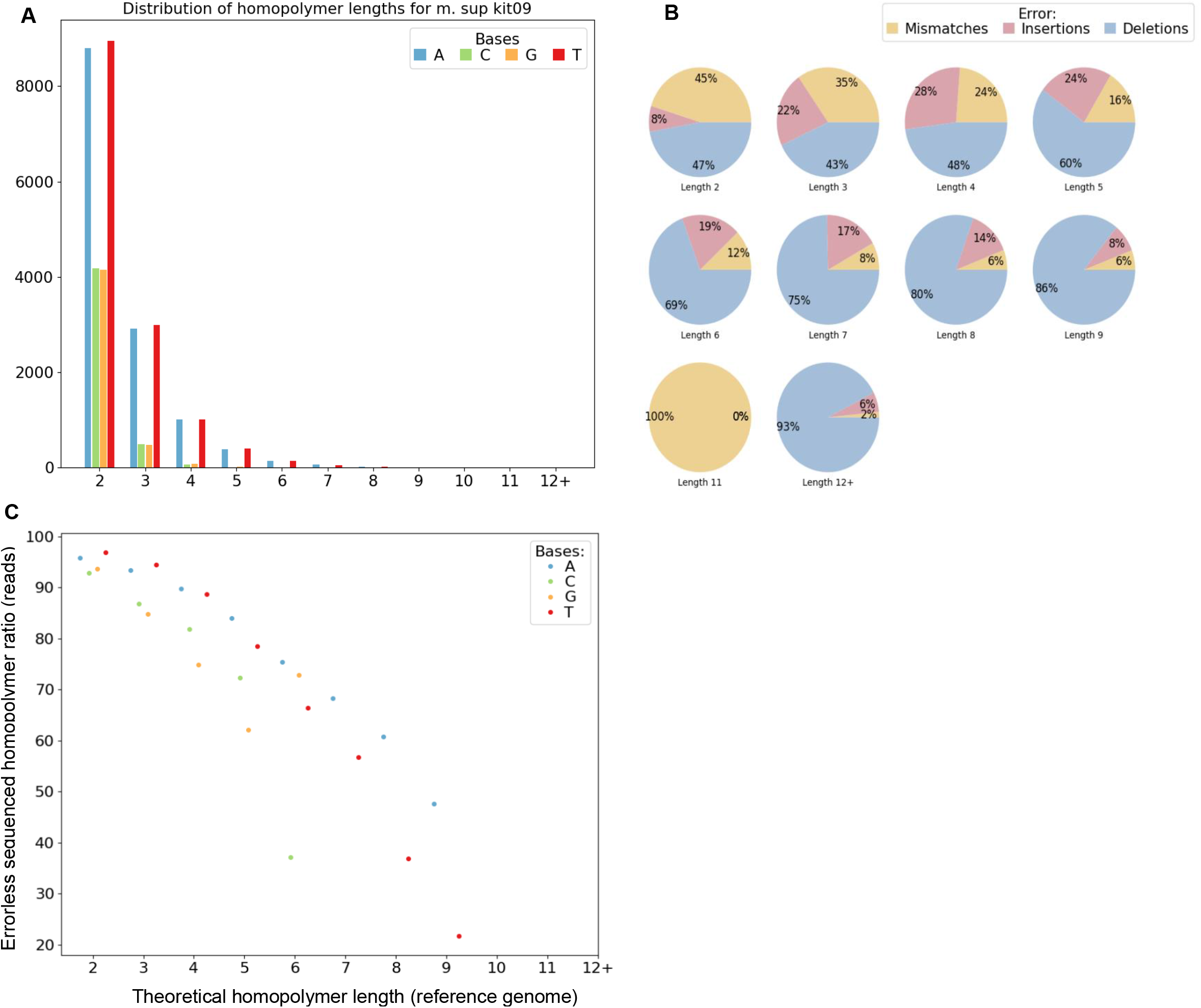
Homopolymer statistics of all reads mapping to MPXV as evaluated by SeqFailR. **A:** Overall distribution of homopolymer classes and lengths for MPXV, **B:** sequencing error type with respect to homopolymer length, **C:** percentage of homopolymers that were sequenced correctly plotted against the theoretical length of the homopolymer

Metagenomic composition was queried throughout the run using Centrifuge. In fact, MPXV could be detected among the first 4000 reads that were generated. After re-basecalling, 99.6% of reads mapped to chordate sequences, and the remaining reads mostly to the MPXV reference (Fig. 3, Tab. 1).

**Fig. 3:**
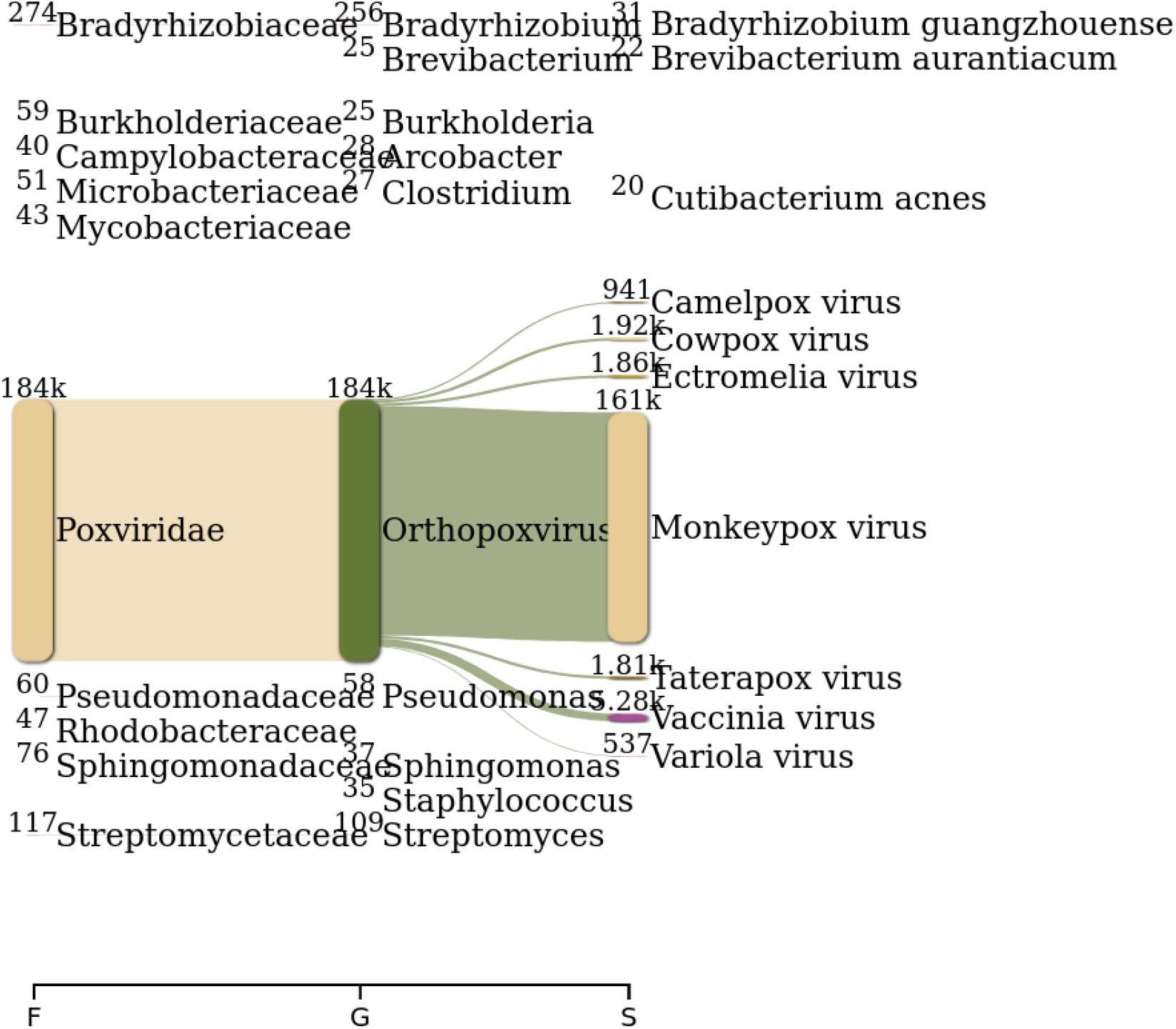
Sankey plot of metagenomic reads after exclusion of reads below the quality threshold and exclusion of human reads

**Tab. 1:** Read distribution of all reads that passed the quality filter after 72h as evaluated by Centrifuge (h+p+v+c index)

| Show | 10 | entries | Search: |  |  |  |  |  |  |  |
| --- | --- | --- | --- | --- | --- | --- | --- | --- | --- | --- |
| Name | Number of raw reads | Classified reads | Chordate reads | Artificial reads | Unclassified reads | Microbial reads | Bacterial reads | Viral reads | Fungal reads | Protozoan reads |
| read_classifications.kit09 | 49,885,777 | 100% | 99.6% | 0% | 0.0331% | 0.373% | 0.00419% | 0.368% | 0% | 0% |

Adaptive sampling was active on half of the flow cell for the entire duration of the run. The rationale was to deplete reads originating from human DNA and thereby to passively increase coverage on the MPXV target. The N50 of an adaptively sampled read was evaluated at 744 bp by MinKNOW. Compared to the other half of the flowcell, the enrichment via adaptive sampling was approximately 2X for reads mapping to the MPXV reference (Fig. 4).

**Fig. 4:**
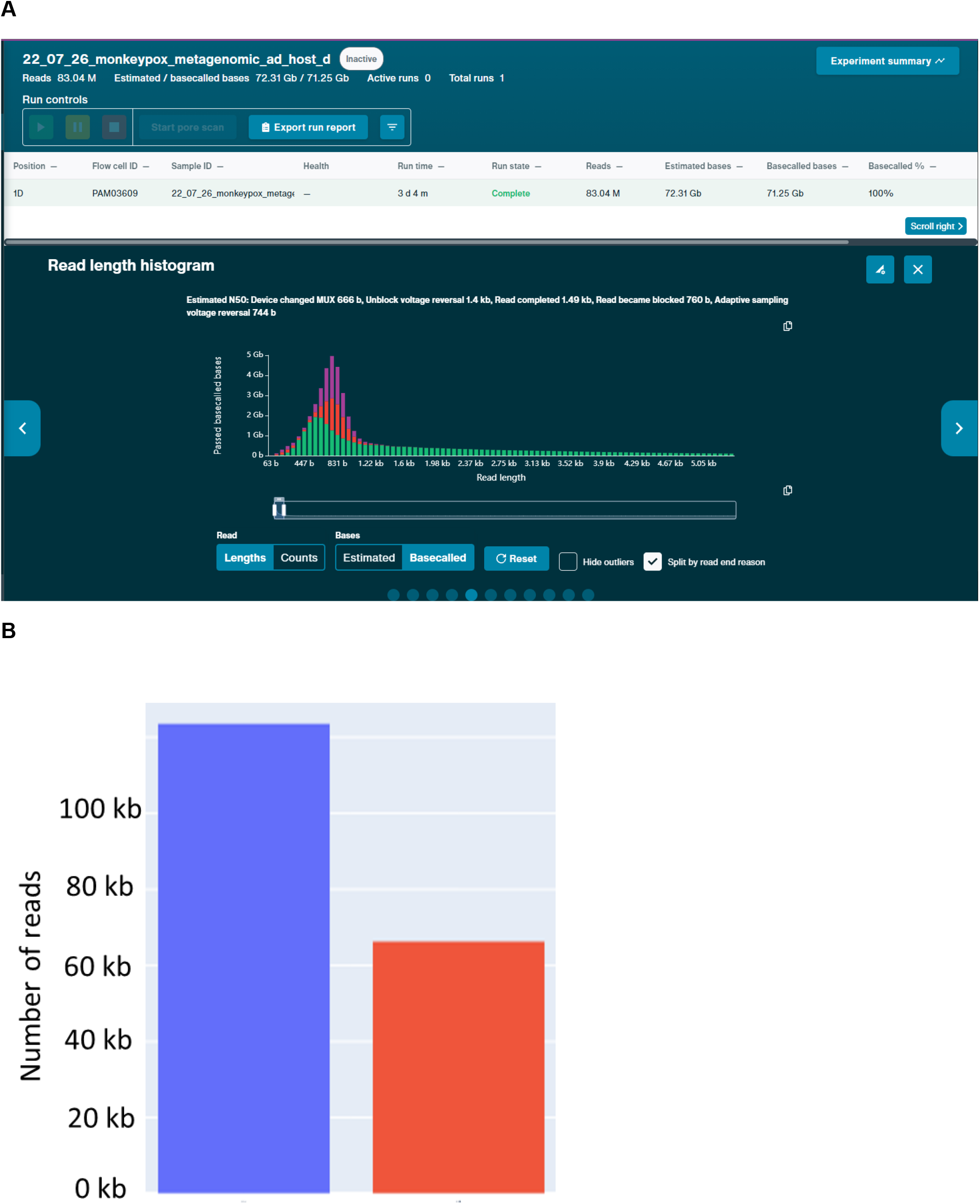
Effect of adaptive sampling decisions on half of the PromethION flow cell A: Read end decision as evaluated by MinKNOW, B: Number of reads mapping against the MPVX reference split by flow cell half (blue: adaptive sampling = ON, red =adaptive sampling = OFF)

Lineage assignment was performed by NextClade and revealed that the MPXV sample belongs to lineage B.1 and Clade IIb frequently observed during the 2022 outbreak. After review of the draft sequence and exclusion of unexpected frameshifts, three notable nucleotide substitutions remained: C101074T, G138682A, C189167T, of which only G138682A leads to an amino acid substitution OPG156:S243F. Moreover, in terms of structural variants, there are two larger insertions located in intergenic regions of the inverted terminal repeats within the MPXV genome (Tab. 2, 3).

**Tab. 2:**
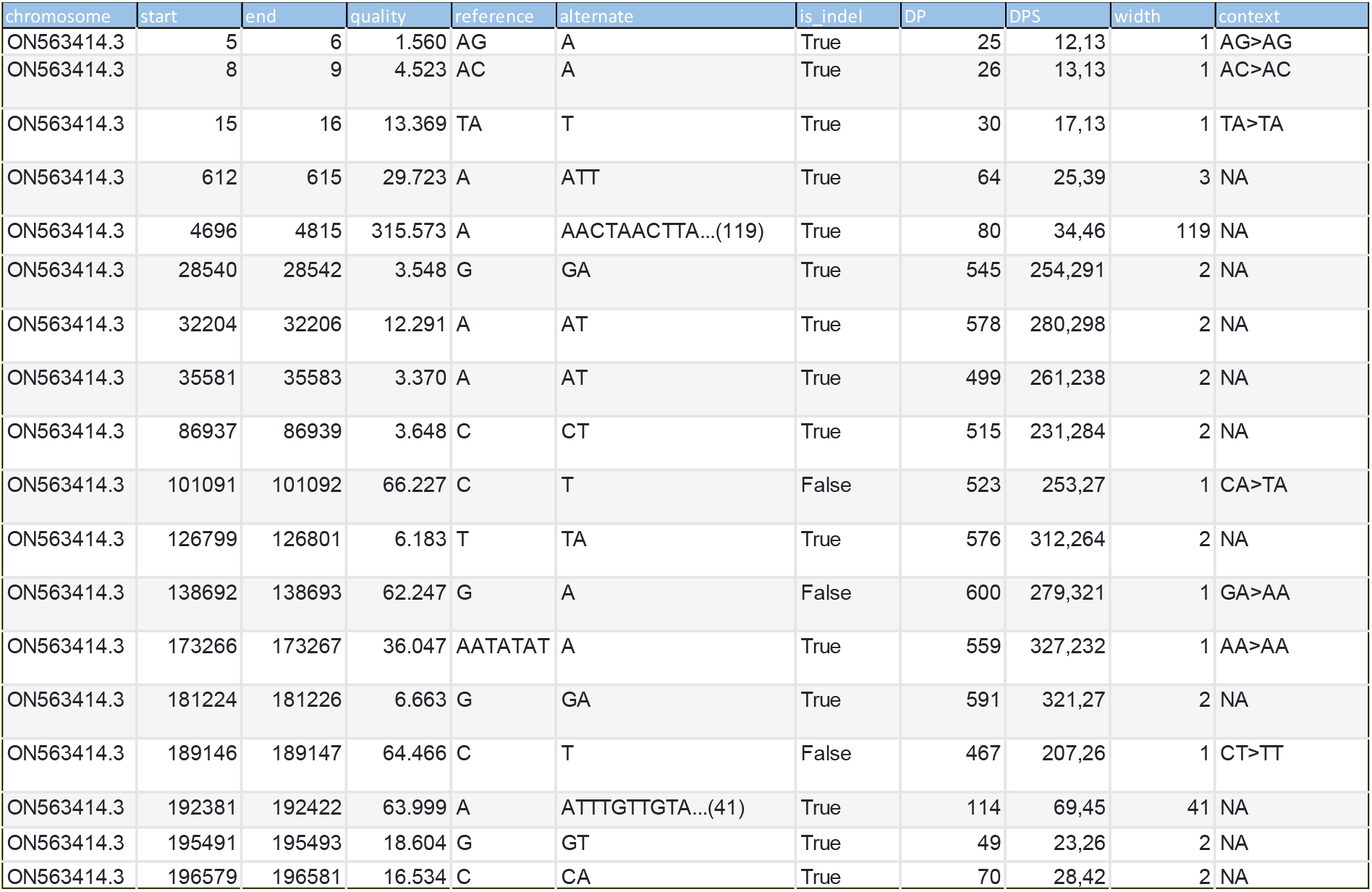
All Variants with respect to ON562414.3 as initially reported by the wf-mpx pipeline

**Tab. 3:** NextClade result for the preliminary draft consensus

| # | i | Sequence name | QC | Clade | Outbreak | Lineage | Mut. | non-ACGTN | Ns | Cov. | Gaps | Ins. | FS | SC |
| --- | --- | --- | --- | --- | --- | --- | --- | --- | --- | --- | --- | --- | --- | --- |
| ? | ? | ? | ? | ? | ? | ? | ? | ? | ? | ? | ? | ? | ? | ? |
| 0 | 0 | ON563414.3 Monkeypox virus isolate M | N M P C F S | IIb | hMPXV-1 | B.1 | 3 | 0 | 0 | 100.0% | 50 | 205 | 4 (4) | 0 |

## Discussion

The present study seeked to characterize the performance of metagenomic nanopore sequencing on a skin lesion sample from a monkeypox case. Human host depletion via adaptive sampling led to a successful, albeit moderate, enrichment of ca. 2X for the MPXV fraction. Metagenomic composition revealed that this enrichment of MPXV sequences was achieved despite a massive excess of human DNA (>99,5%) exceeding previously reported ratios (Marquet et al. 2022). In general, depletion should result in a significantly more favorable human host to pathogen nucleic acid ratio, which would of course increase the taxonomical resolution of microorganism and speed up clinical management of acute infections. A variety of ONT’s adaptive sampling strategies have been previously demonstrated (Payne et al. 2021) and could be applied for human host depletion in metagenomic studies (Marquet et al. 2022, Cheng et al. 2022). Working with clinical samples characterized by high amounts of host DNA, Marquet et al (2022) demonstrated a 1.70-fold increase in total sequencing depth after human host depletion. Cheng et al (2022) could increase microbial sequence yield at least 8-fold while adapting human host depletion in another clinical setting. In principle, the method works well, but is not without limitations, such as increased pore blockage. Another known issue of human host depletion via ONT’s adaptive sampling option in MinKNOW is the usage of its general-purpose alignment program minimap2 (Li et al. 2018). Previous studies showed that around 25% of human reads could not be rejected accurately by this software, thus wasting sequencing resources (Payne et al. 2021, Marquet et al. 2022). Addressing this problem, ReadBouncer was introduced by Ulreich et al. (2022) as an improved classification tool. It shows a higher read sensitivity than other state-of-the-art classification tools for adaptive sampling, while retaining a high specificity. A different potential limitation is the fixed target reference, i.e., the adaptive sampling reference is chosen at the beginning and stays constant during the run. The tool BOSS-RUNS, a further development in this field, can generate dynamically updated decision strategies where coverage is tracked throughout the run and the adaptive sampling target list is kept updated based on the coverage resp. species abundance (Weilguny et al. 2023). Lastly, adaptive sampling is also reliant on read length and increasing the read length will improve abundance of target species (Martin et al. 2022). Both approaches, improvements in the laboratory workflow leading to longer reads, as well as the afore mentioned and upcoming bioinformatic adaptations will make adaptive sampling even more attractive in the future.

One of the complications that afflict a subset of monkeypox patients is bacterial superinfection of the skin lesions (de Sousa et al. 2022). Metagenomic sampling with real-time species determination has the advantage to give an overview of bacterial and viral sample composition within minutes after sequencing start, thus either confirming or refuting the presence of the target pathogen and/or additional pathogens. In this study, we could confirm MPXV as the predominant pathogen in data generated already 10 minutes after sequencing started.

Unlike RNA viruses, the double-stranded DNA of orthopoxviruses is very stable and their DNA polymerase has proofreading exonuclease activity, resulting in a low mutation rate. MPXV of clade IIb most probably originated from clade II in a 2017 outbreak in Nigeria (Isidro et al. 2022). Phylogenetically, these two clades differ by a mean of 50 SNPs, which is roughly 6 to 12-fold more than previous estimations of the substitution rate for Orthopoxviruses (1-2 substitutions per genome per year) (Firth et al. 2010, Isidro et al. 2022). In the present study, we can confirm that the underlying MPXV strain belongs to clade IIb and thus confirm the already known genome sequence differences between clade II and IIb (Isidro et al. 2022). Clade IIb exhibits molecular signatures that can been associated with the potential action of apolipoprotein B mRNA-editing catalytic polypeptide-like 3 (APOBEC3) enzymes via viral genome editing (Pecori et al. 2022). Human APOBEC3 proteins introduce GA > AA and TC > TT replacements into viral genomes and are thus part of the cellular defense mechanisms (Sadeghpour et al. 2021). Mutation patterns of this type are indicative of a large amount of human-to-human transmission. With respect to the inverted terminal repeat (ITR) regions at the ends of the MPXV genomic sequence, it has been previously reported that only long read nanopore reads were able to cover them completely (Vandenbogaert et al. 2022). One key question is how the variations mentioned above affect the MPXV transmissibility, pathogenicity, and adaption to the human host.

Conclusively, metagenomic sequencing of a MPXV sample via adaptive sampling on a PromethION-type flow cell allows coverage of the target species to be increased in addition to rapid, real-time metagenomic classification and lineage determination.

